# Physical Interactions Drive Collective Thermoregulatory Behavior in Honey Bees

**DOI:** 10.1101/2024.12.14.628519

**Authors:** Casey E. Lambert, Kyara Vazquez, Zachary P. Nelson, Chelsea N. Cook

**Affiliations:** Biological Sciences, Marquette University, Milwaukee WI USA

**Keywords:** Interactions, Density, Probability, Fanning, Collective Behavior, Honey Bees

## Abstract

Social animals can coordinate complex behaviors, which can affect massive change on the environment. Within groups, individuals can sense the environment and communicate that information with others. Direct contact, like physical touch, is a key method of communication among social animals, and may be a mechanism to facilitate the coordination of collective behaviors that buffer environmental change. Here, we use a collective thermoregulatory fanning behavior in honey bees (*Apis mellifera*) to test the hypothesis that direct physical contact is necessary to perform this behavior. By modulating their ability to engage in physical contact, we establish that honey bee workers must touch each other to coordinate the fanning response. We then manipulated social density by changing the physical space the bees occupied to modulate likelihood of contact and found that in high social densities, honey bees are more likely to fan. Using video tracking, we then verified that bees in higher social density indeed have more direct physical interactions. This work identifies a mechanism of communication and potentially information synthesis in an ecologically relevant collective behavior. By understanding the ways in which animals communicate, we may be able to pinpoint the mechanisms of resilience that social insects evolved to manage a changing world.

## INTRODUCTION

Social organisms can affect massive change on their environment through coordinated collective behaviors. Locust marches clear croplands as they migrate [1], termite mounds manage their own internal environment as well as shape savannah ecosystems [2,3], and mass animal migrations cycle nutrients across the globe [4]. Collective behavior emerges from coordination of many individuals, potentially providing resiliency to a changing environment [5,6]. Coordination occurs when entities sense shifts in local information from others and the environment, and adjust their own behavior accordingly [7]. Information can be indirect, such as when schooling fish simply watching others move in a particular direction and move with them [8], or it can be direct, such as a honey bee performing a waggle dance to share the location of a food source [7,9,10]. The perception and use of information, especially in regards to collective action, has broad appeal in many fields and applications such as human cultural evolution [11], information and cyber security [12], and social psychology [13]. Defining mechanisms of information use provides important insights into how individuals and systems may maintain resiliency in a changing environment.

Social insect societies engage in many mechanisms of communication, enabling them to manage rapidly shifting environments, and makes them an excellent model system. One major mechanism of communication is direct physical contact. Direct physical contact requires physical touch, often occurring with sensory appendages like antennae, mouthparts, or legs, and can establish and maintain social structures that are the hallmarks of insect societies. Direct physical contact can comprise aggressive interactions that establish dominance hierarchies and reproductive division of labor, as in many species of *Polistes* wasps [14–16]. In honey bees, direct physical contact may promote reproductive drone comb building when the colony reaches a critical mass of 4000 workers [17]. When recruiting nestmates to a new nest site, ants engage in tandem running, which occurs when a recruited ant follows a leader ant by touching the leader’s hind legs with its antennae [18]. Direct physical contact can also coordinate worker division of labor by initiating tasks, such as when inactive foragers begin to forage after an increase in interactions with successful foragers, which also occurs if contact occurs with glass beads that smell like foragers [19] and when paper wasps (*Polybia occidentalis*) bite nestmates to stimulate foraging activity after foragers are removed [20]. In many of these cases, the environment has changed, and information is being communicated to be able to manage that change, although what exactly is being communicated isn’t often known. Direct communication is therefore a likely mechanism driving the coordination of some collective behaviors in response to environmental conditions.

Regulating shifts in temperatures via thermoregulation is among the most critical tasks for social insect societies [21]. Honey bees are cavity nesters that rear sedentary offspring [22]. As such, adults must work together to manage the temperature inside the colony. Brood must be kept at 35°C; development at higher temperatures may result in impaired learning as adults, physical malformations, or death [23–25]. To manage colony temperatures, honey bees perform several thermoregulatory behaviors, such as evaporative cooling via spreading water [26], bearding [27], heat shielding [28] and fanning [29,30]. Fanning occurs throughout the colony as well as at the entrance to circulate air, bringing hot air out and allowing cool air to infiltrate. Fanning is modulated by both thermal and social environment: honey bees are more likely to fan when in groups compared to when as a single individual [31], and larger groups of adult bees fan earlier to combat rapid rates of temperature change [32,33]. Individual adult bees are more likely to fan when able to physically touch a larva, but fanning is reduced when separated from it. Although the social environment modulates the fanning response, the role that social interactions and information use plays in organizing this important thermoregulatory response is unknown.

Here we create specific social environments to test the hypothesis that tactile cues drive the thermoregulatory fanning response in honey bees. First, we identify whether preventing interactions between individuals reduces the fanning response by physically separating individuals. We then examine whether social density modulates the fanning response by changing the size of the physical space they occupy. To understand the social mechanism driving changes in the fanning response, we tracked individuals in high- and low-density groups. From this analysis, we determine that tactile cues, specifically head-to-head interactions drive the fanning response. Our work highlights the importance of information transfer in collective behavioral responses, which are especially important when organizing to manage a changing environment.

## METHODS

### Cage Separation Experiment

#### Husbandry

Ten *Apis mellifera l*. colonies were managed at a bee yard in Boulder, Colorado USA on the University of Colorado’s East campus. Colonies were maintained in 10-frame wooden hive bodies with plastic frames. Supplemental feeding of 2:1 sucrose:water solution occurred as needed. Experiments were conducted 1 May 2014 to 1 September 2014.

#### Identification of Fanners

To identify thermoregulatory fanners, we observed the entrance for bees that were standing still and fanning their wings in place for 10 seconds, with a curved abdomen. Fanners often faced into the colony. The thermoregulatory fanning stance is different from Nasanov fanning (AKA scenting), which also occurs at the entrance of the colony, but can be distinguished from thermoregulatory fanning as their abdomens are straight, angled upward, and a gland at the tip of the abdomen is often exposed [34]. Although fanning occurs throughout the colony, we collected fanning from the entrance of the colony due to their accessibility.

To collect bees, we used forceps to grab the legs of the fanners, then placed them into cages. Cages were cylindrical (r=4.25cm, h= 4cm, total area = 226.98cm^3^) constructed of 0.25in x 0.25in galvanized steel hardware cloth (McGucken Hardware). Five fanning honey bees were collected into 1 of 3 different cage types: open, divided with 1 layer of hardware cloth, divided with 2 layers of hardware cloth. When the bees were placed into the divided cages, an individual could interact with 2 others, functionally creating groups of 3, so we also tested groups of 3 bees in open cages as a control (Fig 1). The treatment group of 3 bees in open cages was a similar experiment being performed in the same time frame, and has a lower sample size (n=13) compared to our groups of 5 open cages (n=61), single mesh divided cages (n=57), and double mesh divided (n=44). Once bees were collected, they were placed into the heating chamber, described below, for a 25-minute room temperature (average = 25.2°C) acclimation period.

**Figure 1:**
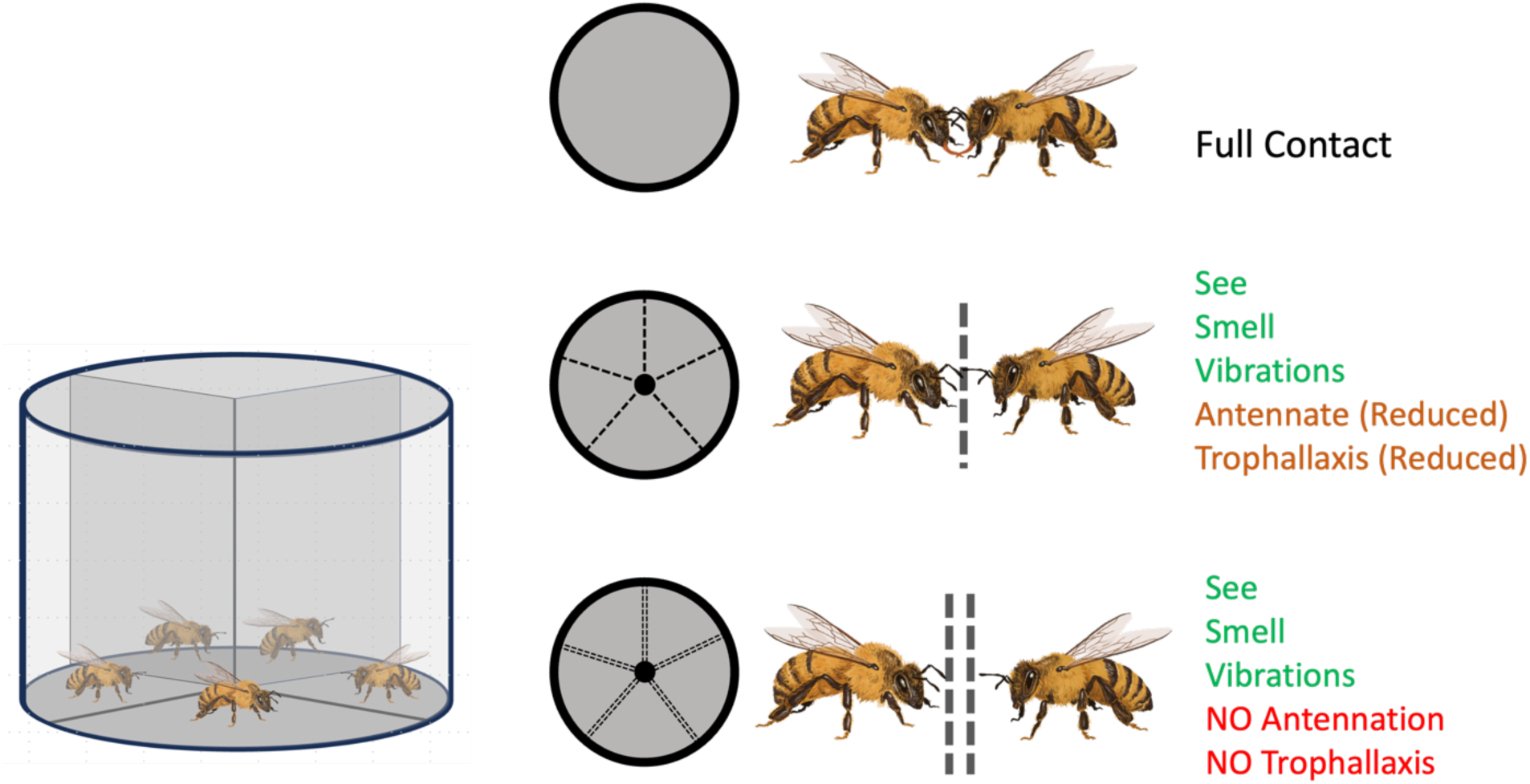
Creating separated cages to reduce interactions. Groups of 5 or 3 bees either moved freely and had full contact, or groups of 5 bees were separated by hardware cloth mesh that reduced or prevented interactions. Bee drawings done by Impact Media Lab.

##### Fanning Assay

To evaluate the fanning response, we heated cages of bees in 1-gallon jars (Uline) on hot plates (Ohaus Guardian 5000). The cages did not touch the glass but were suspended on wooden stands inside the jar. We monitored air temperature with Cole Parmer high accuracy (±0.3°C) digital thermometer. We heated the air temperature inside the jars at 1°C per minute, starting at room temperature. We monitored the cages for fanning behavior, characterized similarly to fanning identification at the hive: flapping wings, curved abdomen, stationary for 10 seconds. We recorded the first instance number of fanners and the temperature that they fanned at. Bees were counted as fanning within a bout if they began within the 30 seconds after the first bee. Assays concluded when all activity ceased.

##### Statistical Analysis

To analyze fanning behavior in the group size and cage separation experiments, we created a four-factor predictor variable which combined the number of bees and the separation of them (i.e.: no separation, 5 or 3 bees; 1 layer, 5 bees; 2 layers, 5 bees). To analyze the probability of fanning, we performed a logistic regression with logit transformation using a generalized linear mixed-effect model on probability of fanning. The response variable for this model was a two-column probability of number of fanners and number that did not fan. The temperature of fanning was analyzed as a generalized linear mixed-effect model with a log transformation (with gaussian family and log link), as the temperature response variable was not normally distributed. To analyze the magnitude of effects of the predictor variable, we performed a type II ANOVA Wald Chi square test. Both included the random effect to account for hive effects. For post-hoc pairwise comparisons we performed Tukey tests. All analyses were performed in R ([35]version 4.2.2) using RStudio ([36]version 2023.06.1+524) packages lme4, car, and emmeans. Graphs were created using ggplot2.

### Density experiment

#### Colony Husbandry

Ten *Apis mellifera l*. colonies were managed on the roof of Wehr Life Sciences on Marquette’s campus in Milwaukee Wisconsin USA. They were maintained in 10-frame Langstroth boxes, and boxes were added as colonies grew through the summer. Frames were plastic (Acorn). Supplemental feeding of 2:1 sucrose:water solution and pollen substitute occurred as needed. Colonies were also treated for mites in August using Apiguard (Mann Lake). All experiments were conducted May 2021-September 2021.

To collect fanners, we identified them using the criteria discussed previously, then used forceps to grab the legs of the bee and placed them into cylindrical cages made of a plywood base, hardware cloth sides, and a plexiglass opening which was a spinning lid to quickly close the cage to prevent escapes. As bees were collected, they were placed directly into two treatment groups: large (r=4.25cm, h= 11.5cm, total area = 652.56cm^3^) or small (r=4.25cm, h= 4cm, total area = 226.98cm^3^) cages. Bees were collected into groups of 10. Cage collection order was alternated each collection bout. 3-4 collection bouts were typical per day. Collections typically lasted less than 5 minutes, however if they took longer, cages were placed in the shade. Cages were returned to the lab and placed directly into the heating chambers to acclimate for 25 minutes.

#### Fanning Assay

To evaluate the fanning response, we heated cages of 10 bees placed inside of 1-gallon jars (Uline) on hot plates (Ohaus Guardian 5000). The cages did not touch the glass but were suspended on wooden stands inside the jar. We monitored air temperature with probes (109, Campbell Scientific) and analog input module (Granite Series Volt 116, Campbell Scientific) using their software Surveyor. We heated the air temperature inside the jars at 1°C per minute, starting at room temperature (average=23.67°C). We monitored the cages for fanning behavior, characterized similarly to fanning identification at the hive: flapping wings, curved abdomen, stationary for 10 seconds. We recorded the number of fanners and the temperature that they fanned at. Often, separate fanning bouts would occur, where bees would begin to fan, stop fanning, then begin at a higher temp. As such, we kept track of these bouts separately as the initial fanning bout and the fanning bout with the maximum number of fanners, where were sometimes the same bout. Bees were counted as fanning within a bout if they began within the 30 seconds after the first bee.

#### Statistical Analysis

To analyze the probability of fanning in the two social densities, we performed a logistic regression with logit transformation using a generalized linear mixed-effect model on probability of fanning. The response variable for this model was a two-column probability of number of fanners and number that did not fan. The temperature of fanning was analyzed as a generalized linear mixed-effect model with a log transformation (with a gaussian distribution and log link), as the temperature response variable was not normally distributed. The predictor variable was social density (low or high). To analyze the magnitude of effects of the predictor variables, we performed a type II ANOVA Wald Chi square test. Both included the random effect to account for the different collection hives. For post-hoc pairwise comparisons we performed Tukey tests. All analyses were performed in R ([35] version 4.2.2) using Rstudio [36], packages lme4, car, and emmeans. Graphs were created using ggplot2.

#### Video Tracking

##### Cages

To evaluate whether the behavioral mechanism that drives the fanning response were physical interactions, we quantified number of interactions and interaction duration. To do this, we video recorded fanning assays in high density and low-density social environments. We created a cage as 2-dimentional as possible to enhance video recording and individual tracking quality. These cages were rectangular, as opposed to cylindrical as compared to our previous behavioral assays. Trial cages were constructed with wooden sides, a glass removable top panel, and an aluminum hardware mesh base which was covered with a silicone mat. Trial containers measured 13.5 cm x 14.5 cm x 2.5 cm. Hardware mesh was used to wall off sections of the cage to alter the housing area and encase the temperature probe. The high-density (smaller) housing was 13.5 cm x 7 cm x 2.5 cm; with a floor space of 94.5 cm^2^ and total volume of 236.25 cm^3^. The low-density (larger) housing was 13.5 cm x 14.5 cm x 2.5 cm but a 3.5 cm x 6.5 cm x 2.5 cm section of was blocked off by the temperature probe for a final floor space of 173 cm^2^ and total volume of 432.5 cm^3^.

##### Collecting Bees

To collect bees, we followed the same protocols as stated above by identifying fanners at the entrance of the colony and using forceps to grab them by their legs. To keep track of individuals during video tracking, we individually marked the bees using different color water-based acrylic paint markers (Montana Brand) as we collected them, before placing them into the two different cage size treatments.

##### Video recording

To ensure high quality videos, we created a filming box that was well-lit but reduced reflections. To do this, we created a recording (39cm x 39cm x 57cm dimension) box, painted the inside flat white, and added two circular (14cm diameter) holes for lights on opposite sides. We used two 8.5 watt (60-watt equivalent) daylight bright LED (GE) to light the box. We created a hole at the top of the box for the video camera to cradle in for recording. We used a video camera (Panasonic HC-V800) with a SD card (SanDisk 128mb microSD) to record. Temperature conditions were monitored using a Campbell Scientific *109 Temperature Probe* wired to a Granite Series *Volt 116* and read and recorded through SURVEYOR version 1.01. Trials were recorded with a high-definition *Panasonic HC-V800* video camera. Once placed in trial housing, the cages were placed in video recording boxes and allowed to acclimate for 25 minutes. Temperature conditions increased using a hot plate at an average rate of 1°C/min, and all trials were conducted under standardized light conditions. We stored videos locally and backed up on cloud services provided by Marquette University IT through Microsoft Sharepoint.

##### Video Tracking

We used a software program called ABCTracker [37] to track individual location and interactions. ABCTracker identifies and tracks the body and head of individuals as well as planar coordinates associated with time, which allows for overlap to be identified. Video was tracked at a resolution of thirty frames per second. After running the initial process tracking process, tracks were manually corrected.

##### Tracking Analysis

We compared the interaction dynamics under increasing temperature conditions for groups of ten bees across two cage sizes; high density and low density. Three trials were conducted for each cage size, totaling 6 trials and 60 individual bees.

To ensure comparability across trials, we analyzed the four minutes of activity prior to the start of temperature fanning (disregarding Nasanov fanning). This allows us to account for normal variation in acclimation and initial fanning temperature across groups. An interaction was defined as lasting at least 1 second(s). Interactions between unique pairs of individuals happening less than 1s apart were combined into a single instance. Interactions were capped at 60s to reduce the effect of individual bees resting next to one another [38]. Interactions were categorized as being head-to-head (HH), head-to-body (HB), or body-body (BB). As it was possible for multiple interaction types to happen simultaneously, we set an interaction hierarchy which ranked HH first, then HB, then BB. For example, if both a HH and an HB interaction were registered within the frame, it was classified as HH.

We aggregated individual level data, and analyzed the type, number and duration of interactions across the high- and low-density cages. A series of Shapiro-Wilk tests concluded that the data was not normally distributed (Table S1a and S1b), so we used a Wilcoxon Rank-Sum Test to compare results across the two cage sizes. All analyses for this experiment were performed in R version 4.3.1 [35] and R studio version [36] 2023.06.0+421.

## RESULTS

### Physical separation reduces the fanning response

Separating bees significantly reduced the probability of fanning (Figure 2A: Analysis of Deviance: χ² =85.38, p<0.001). We found that honey bees were significantly more likely to fan when they in open cages, not divided by single (p<0.001) or double mesh (p<0.001). There was no significant difference in the probability of fanning between single or double mesh cages (p=0.42). There was also no significant difference in probability of fanning between groups of 5 bees and groups of 3 freely moving bees that were in open cages (p=0.99).

**Figure 2:**
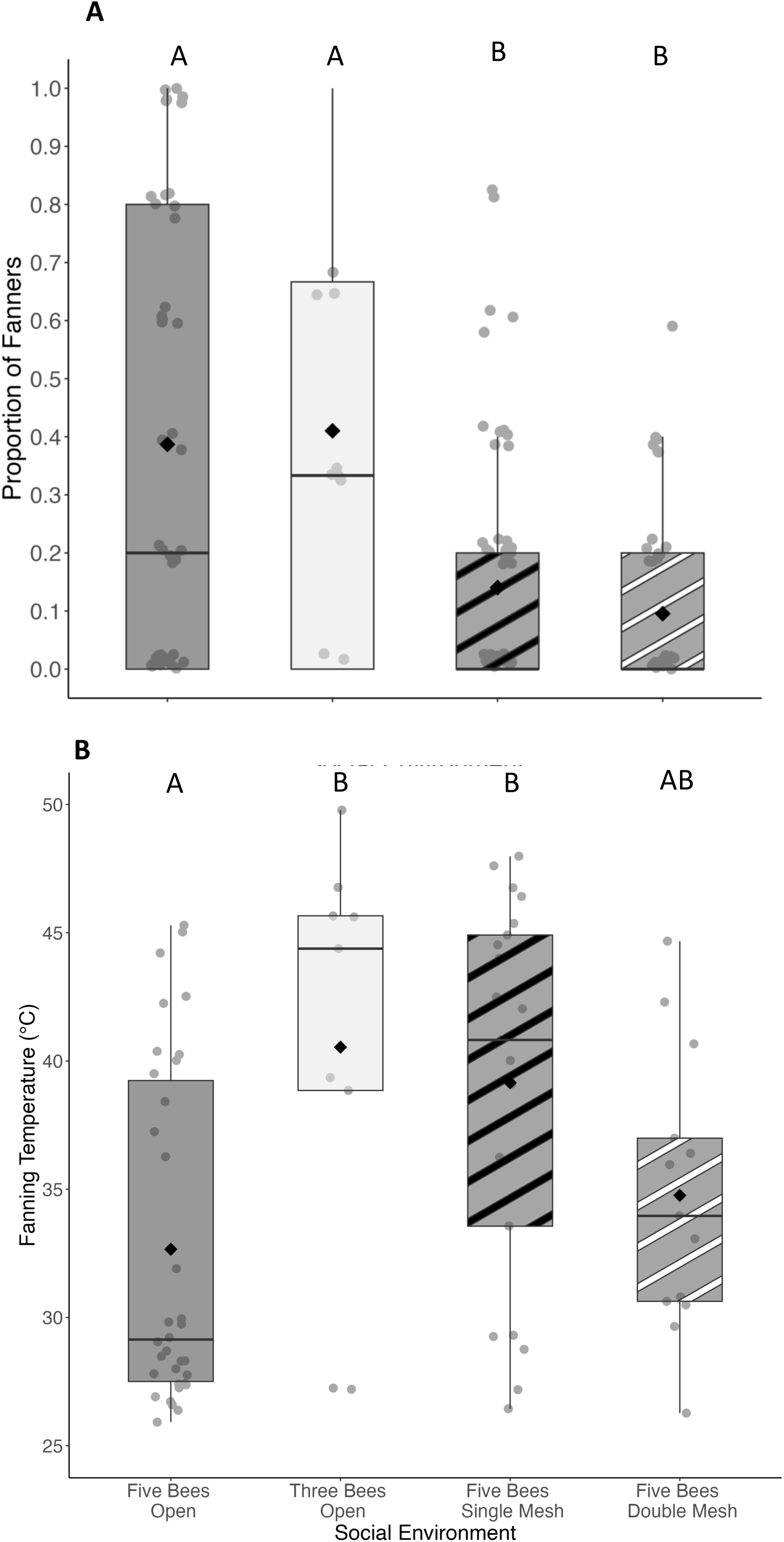
The fanning response of honey bees in varying group sizes and degrees of separation. A) Honey bees that are able to freely interact in groups of 5 (n=61) or 3 (n=13) are significantly more likely to fan compared to when interactions are restricted by a single layer of mesh (n=57) or a double layer of mesh (n=44). B) Honey bees fan at lower temperatures when they are freely moving in groups of 5, and fan at higher temperature when they are in smaller groups of 3 or divided. Letters above boxes correspond to Tukey post-hoc test at p<0.05.

The temperature at which bees began to fan was predicted by both group size and type of separation (Figure 2B: Analysis of Deviance: χ² =15.02, p=0.002). Bees that were freely moving in groups of 5 fanned at an average of 32.7± 1.2°C, which was significantly lower than groups of 3 freely moving bees (mean = 40.5± 2.33°C; p=0.011) and single mesh divided groups of 5 (mean = 38.5± 1.5°C, p=0.014). Bees in double mesh fanned at an average of 34.1± 1.86°C, which was not significantly different than any of the other groups (p>0.05).

### Higher Social Density Increases the Fanning Response

When bees are in the high-density social environment, they are significantly more likely to fan, compared to the lower density environment (Figure 2). There were significantly more bees in the first fanning bout (Fig 2a; Analysis of Deviance: χ² =8.62, p=0.003;), as well as the bout with the most fanners during the assay (Fig 2b; Analysis of Deviance: χ² =7.44, p=0.006). There was no significant difference in the temperature that the fanning bees initiated the fanning response (Fig 3a; Analysis of Deviance: χ² =2.17, p=0.14) or the temperature that bees were fanning when the highest number of fanners were fanning (Figure 3b; Analysis of Deviance: χ² =0.41, p=0.52).

**Figure 3:**
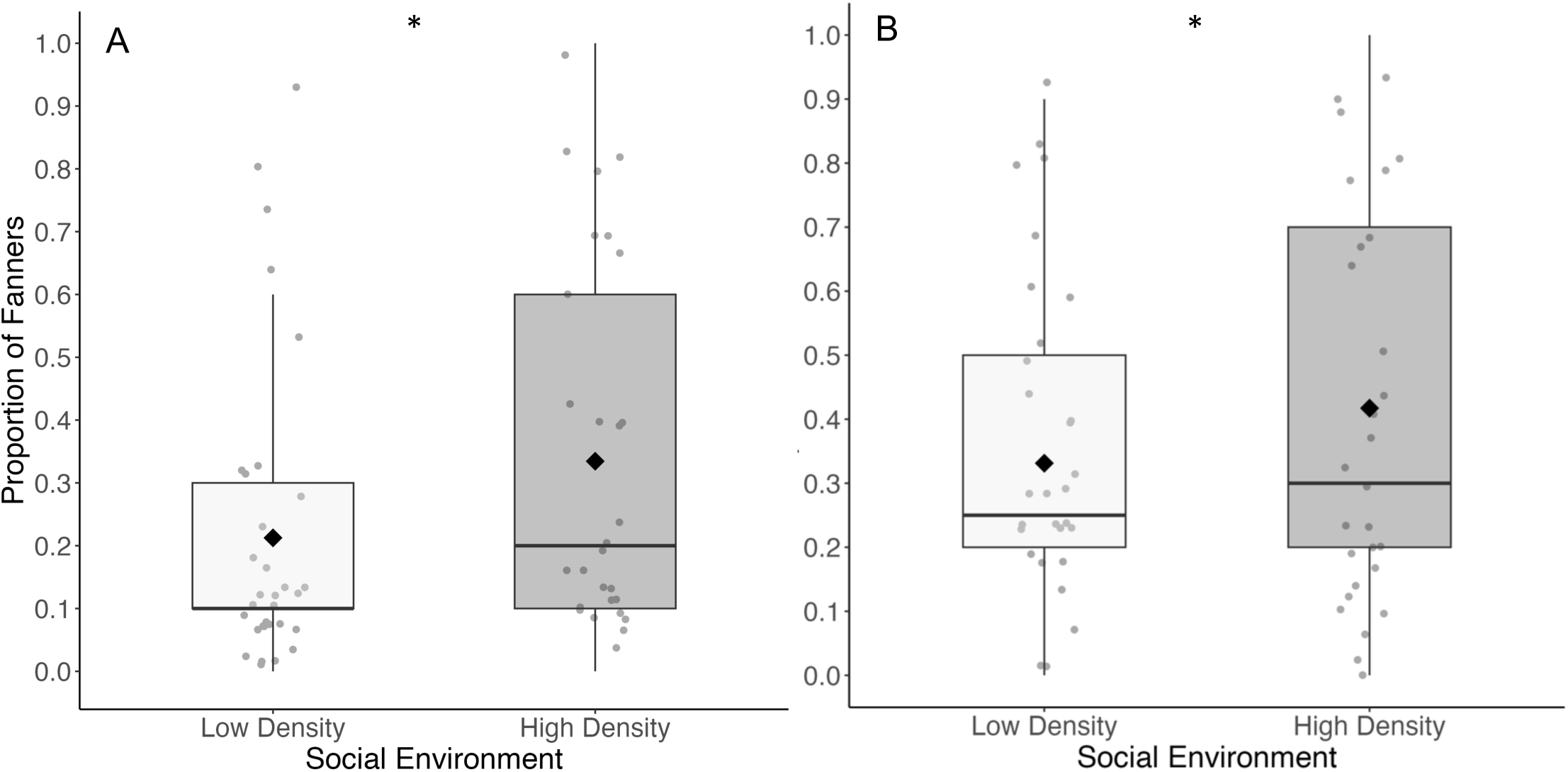
Honey bees are more likely to fan when in high density social environments compared to lower density environments. This includes both the initial fanning bout (A) as well as the fanning bout where the greatest number of bees fan (B). Number of small cage trials = 29, number of large cage trials = 32. Dots are the proportion of fanners in a given fanning bout (number of fanners/10). Boxes represent the interquartile range (IQR) and whiskers represent 1.5*IQR. Thick bars are medians and diamonds are means of all the data. Asterisks indicate significance according to logistic regression at p < 0.05.

#### Higher density social environments and interactions

The social environment did not significantly impact the type of interactions that the bees were engaging in. Overwhelmingly, bees had more head-to-head (HH) interactions (Figure 4), accounting for 71.7% of interactions in the low-density cages, and 80.1 % of interactions in the high-density cages. Bees in low-density cages engaged in 27% of head-to-body (HB) interactions and 1.6% body-to-body (BB) interactions, compared to 10% head-to-body and 9.3% BB interactions in high density cages. Low-density cages engaged in 1712 interactions total and high-density cages engaged in 4098 interactions total (Table S2).

**Figure 4:**
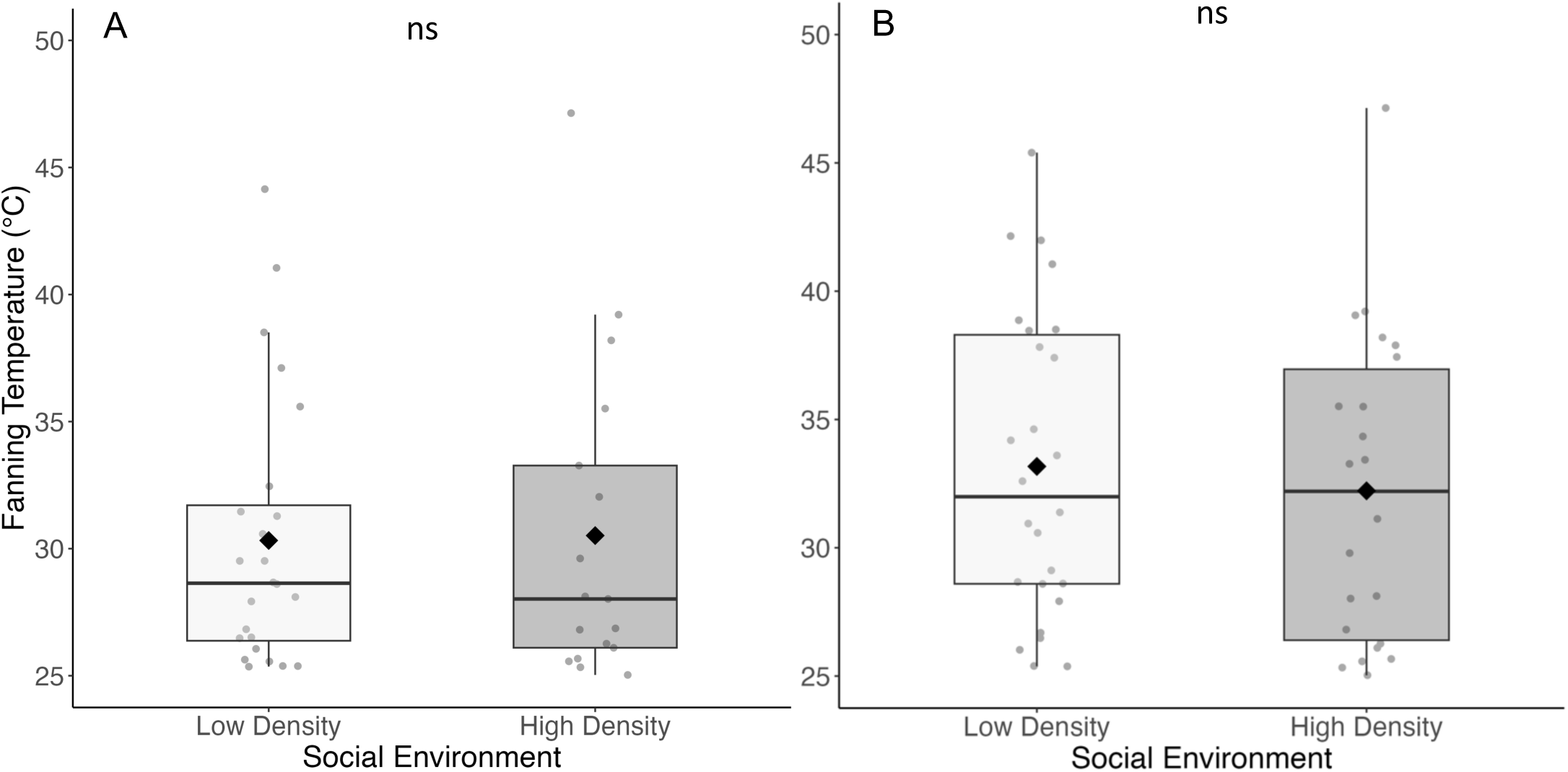
Fanning honey bees collectively fan at the same temperature, regardless of the density of their social environments. A) Honey bees begin to fan at statistically similar temperatures and B) the temperature at which the maximum number of bees are fanning is statistically similar when bees are in high and low density social environments. N=27 for both groups where fanning was observed. Boxes represent the interquartile range (IQR) and whiskers represent 1.5*IQR. Thick bars are medians and diamonds are means of all the data.

To further explore how the bees interacted with each other, we quantified the number of interactions each individual was having. In the high-density environments, individuals were having an average of 157 interactions in the 4-minute segment before fanning occurred, which was significantly more than the low-density environment where they had an average of 57.1 interactions (Table S3; Figure 5A; Wilcoxon test: p < 0.001). Honey bees in high-density social environments had significantly more HH interactions than bees in low-density social environments (Figure 5B; Wilcox test: p < 0.001). Bees in high density environments also had significantly fewer HB interactions compared to bees interacting in the low-density environments (Figure 5C; Wilcox test: p = 0.02). There was no difference in BB interactions in the different social densities (Wilcox test: p > 0.05). Full results in Table S4.

**Figure 5:**
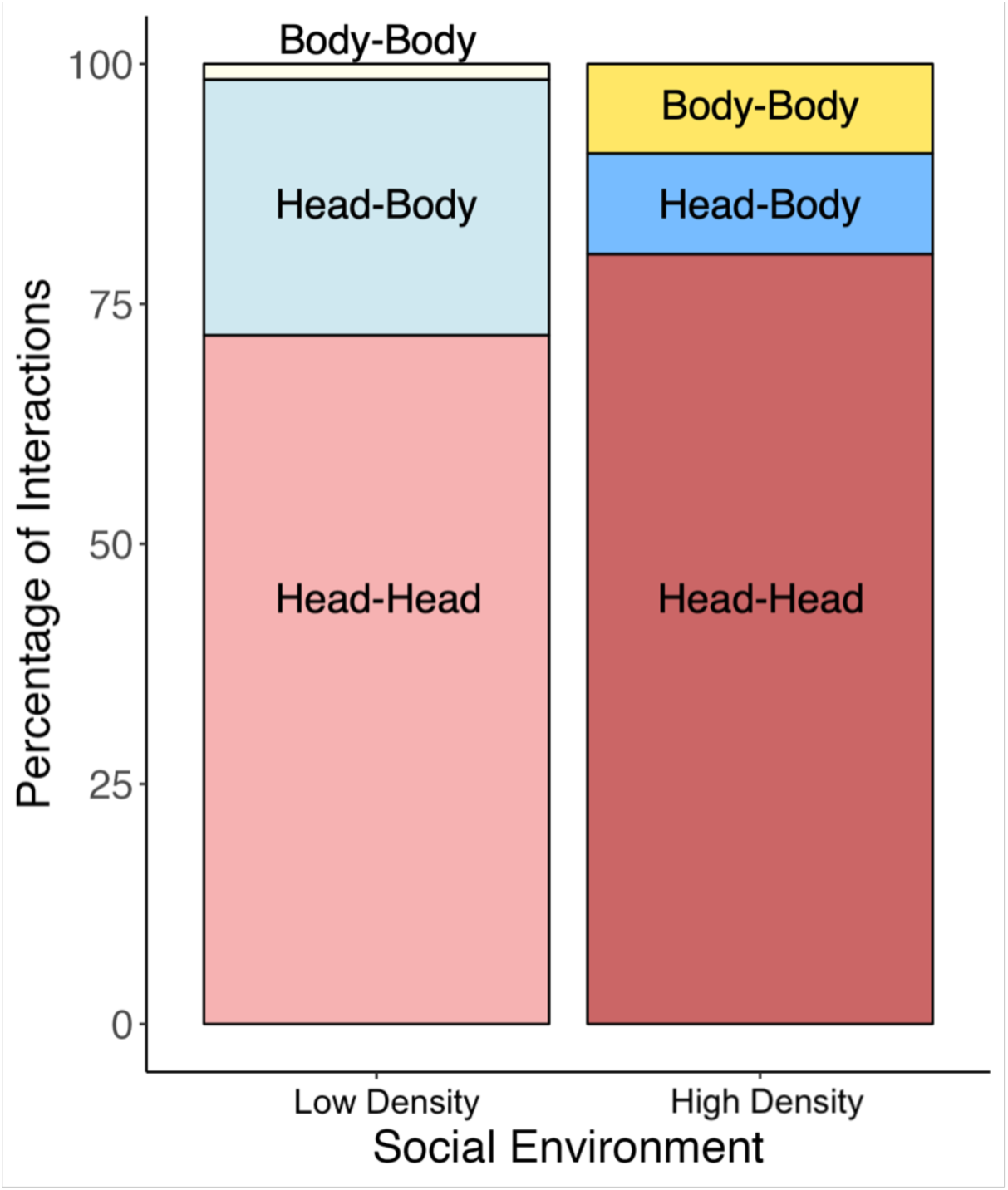
Interacting honey bees prioritized head-to-head interactions over other types of interactions, regardless of social density. Bees interacting in low density social environments, bees had 1228/1712 head-to-head interactions and bees in high density environments had 3286/4098 head-to-head interactions, which comprise about 75% of all interactions in both social environments. Interacting bees performed more head-to-body interactions (456/1712) and very few body-to-body interactions (28/1712) compared to head-to-body interactions (430/4098) and body-to-body interactions (382/4098) in high social density environments.

Duration of interactions did not differ across high and low social densities (Figure 4A; Wilcoxon test p=0.83, n=30 in each group). However, when broken down into specific interactions, there was a significant difference in HB interactions across the two groups (Figure 4C; p<0.001, n=30 in low density, n=10 in high density) which was likely caused from the skewed distribution resulting from the 20 bees across videos having zero head-to-body interactions, attributed to clustering and head-to-head interactions being prioritized (Table S5, S6). There was no difference in the average HH (Figure 4B; p = 0.51, n=30 in both groups), or BB interactions (Figure 4D; p=0.86, n= 12 in low density, n=15 in high density). Full Wilcoxon results in Table S7.

## DISCUSSION

Here, we tested the hypothesis that direct physical contact was necessary for the thermoregulatory fanning response in honey bees. Our results indicate that indeed physical interactions are necessary for the performance of fanning. When bees are prevented from interacting with each other with a barrier, they are significantly less likely to fan (Figure 1). This is further supported by our findings that when the same number of bees are in a smaller space, they are significantly more likely to fan compared to when they occupy a larger space (Figure 2). Finally, we confirm that honey bees in higher social density have more head-to-head interactions compared to bees in lower social density (Figure 5) which correlates with the observed increase in fanning. Together, this work suggests that direct physical contact enables honey bees to coordinate the fanning response.

Honey bees utilize a myriad of modes to communicate, including tactile [39,40], vibrational [41], and pheromonal cues [42]. Here, we hypothesized that physical contact is the most important for fanning behavior. We tested this hypothesis by preventing specific interactions from occurring. The single mesh divisions allowed visual cues, perception of vibration, perception of volatile pheromones, but did reduce tactile cues, such as trophallaxis and antennation. The double mesh divisions allowed visual cues, perception of vibration, and perception of volatile pheromones, but eliminated physical contact. We predicted that when honeybees are prevented from touching, they would fan less. These methods replicate similar experiments that systematically rule out mechanisms of communication. Physical contact between older honey bees inhibited the physiological and behavioral development of younger bees, when they were reared in isolation [43] or separated by a double or single layer of mesh, nurses had forager-like physiology and began to forage earlier [44]. Dor et al. [45] separated worker honey bees with single or double layers of mesh and compared them to isolated bees or freely interacting pairs to identify how dominance between individuals may be established. When in pairs, one bee always developed larger ovaries regardless of separation type, suggesting pheromones do play a role in establishing dominance. However, the bee that developed larger ovaries always had smaller ovaries compared to the dominant bee in the unrestricted pair, indicating that full physical interactions also mattered. Our results align with these previous findings that when bees are prevented from fully physically interacting, they are significantly less likely to fan, which indicates that physical contact is necessary to initiate fanning and adds to the evidence of the importance tactile information transfer in collective behavior.

We show that density also plays a role in a collective thermoregulatory behavior in honey bees, further supporting our hypothesis. We placed a group of 10 bees into a high-density or low-density social environment by changing the size of the testing arena. We found that groups of 10 worker honey bees were significantly more likely to begin to fan when in a high-density social environment compared to a low density one (Figure 2). Bees in high density were twice as likely to begin to fan than in low density. There was no significant difference in the temperature at which the group began fanning (Figure 3). These results reinforce density-dependent interactions as a mechanism of behavioral shifts that play a role in the division of labor in social insects. For example, inactive red harvester ants (*Pogonomyrmex barbatus*) begin to forage when interactions with patrollers hit a specific threshold of 1 ant every 10 seconds or less, which was confirmed by changing the rate of patroller return by introducing glass beads covered in patroller cuticular hydrocarbons [19]. Although our results provide evidence that physical contact mediates the fanning response, we cannot rule out that there is a tactile pheromone or other chemical cue that is also transmitted during touch, similar to the cuticular hydrocarbons of foragers in harvester ants. Physical interactions seem to be providing information about the environment, such as food availability or temperature, and honey bee fanning can inform theories about what and how information is transmitted.

Increased social density leads to increased physical interactions, which likely facilitated the increase in the observed fanning response. In our assays, honey bees interacted extensively, and they prioritized head-to-head interactions, regardless of social density (Figure 4). Our findings show that honey bees have more interactions overall compared to when they are in less dense social environments (Figure 5). Less dense environments lead to longer durations when interacting, which does not correlate with fanning (Figure 6). During head-to-head interactions, eusocial insects engage in antennation and trophallaxis, which are hypothesized to convey important information about environmental conditions [38,46,47]. For example, honey bees can learn odor information when rewarded with the touch of a nestmate, similar to when they are rewarded with food [48]. The number of interactions, and the information they carry, influence whether social insects like honey bees will engage in behaviors, and likely help them coordinate collective behaviors like thermoregulation.

**Figure 6:**
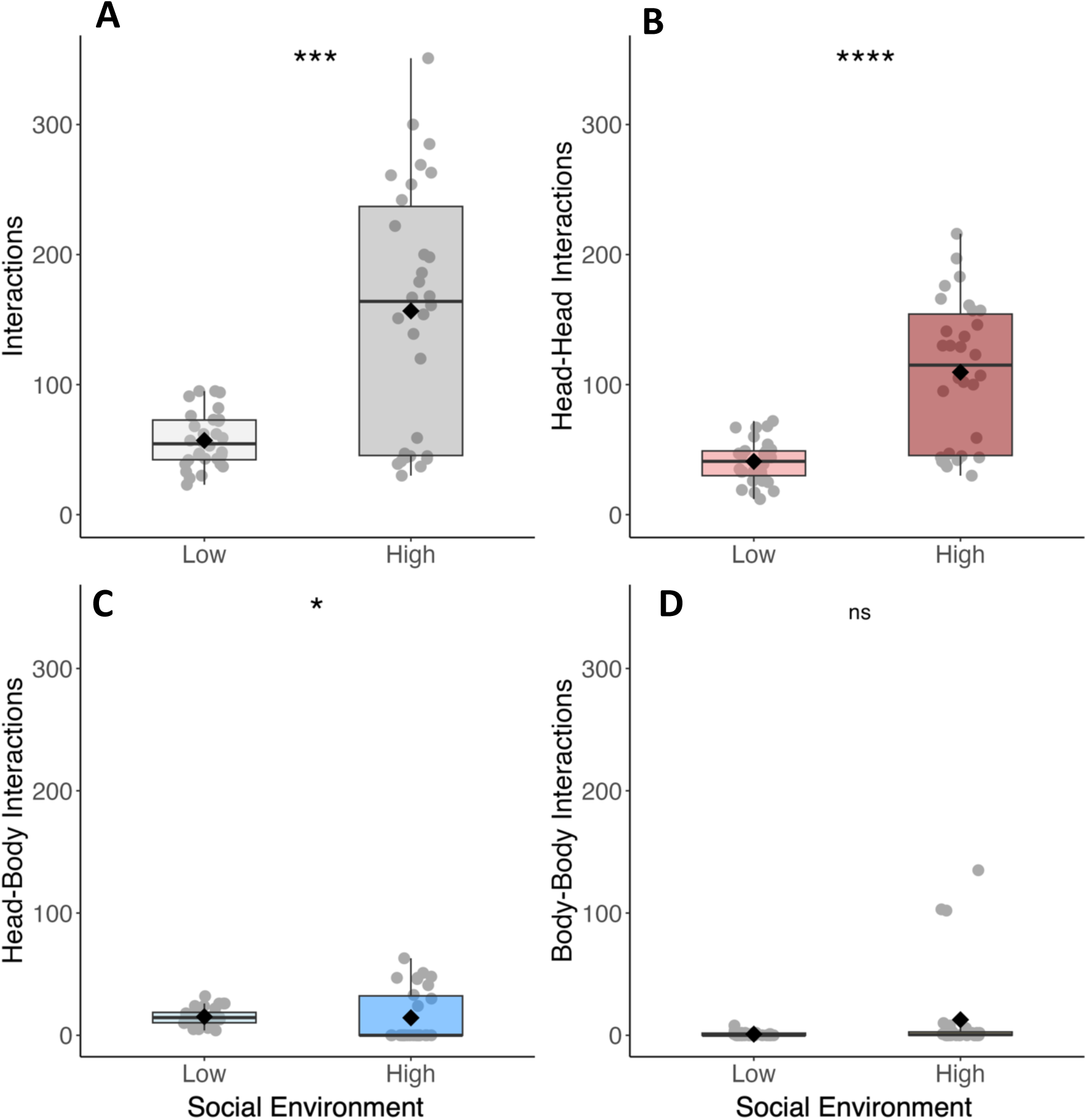
Individual bees in high density environments had more head-to-head interactions. When ten bees were in smaller, high-density cages, they had more A) overall interactions and B) head-to-head interactions. There was no difference in the C) head-to-body interactions or the D) body-body interactions. Interactions are characterized as a boundary around the body of the individual bee overlapping. Head-to-head interactions occur when a diamond shaped boundary over the head of the bee overlap with another diamond shaped boundary with another bee. Interactions are ranked: When both occur at the same time, head-to-head interactions are prioritized. Each dot is the raw number of interactions that an individual engages in during the 4 minutes before fanning. Boxes represent 25-75 quartiles, whiskers indicate 95% of the data. Thick bars are medians and diamonds are means of all the data. Asterisks indicate significance according to Wilcox test at p < 0.05.

Recent hypotheses identify network information theory as a mechanism of maintaining resilience in a complex systems [49], especially in social animal behavior [50]. Complex behaviors like honey bee fanning likely require multiple information inputs, such as temperature of the environment and social environment. Acquired information is potentially synthesized among individuals when they interact to confirm that information, which may help to coordinate subsequent behaviors [10]. In previous work, we found that honey bees are more likely to fan when in groups of 10 compared to smaller groups of 3 or when isolated [31]. In these larger groups, honey bees appear to anticipate the rate of temperature change, as they begin fanning earlier when experiencing a fast rate of temperature change [32]. As such, individual honey bees may be sensing the environment, but need to communicate with others to properly synthesize the information to initiate a collective response. There is still much work to be done to understand what information may be being communicated or how information is shared.

**Figure 7:**
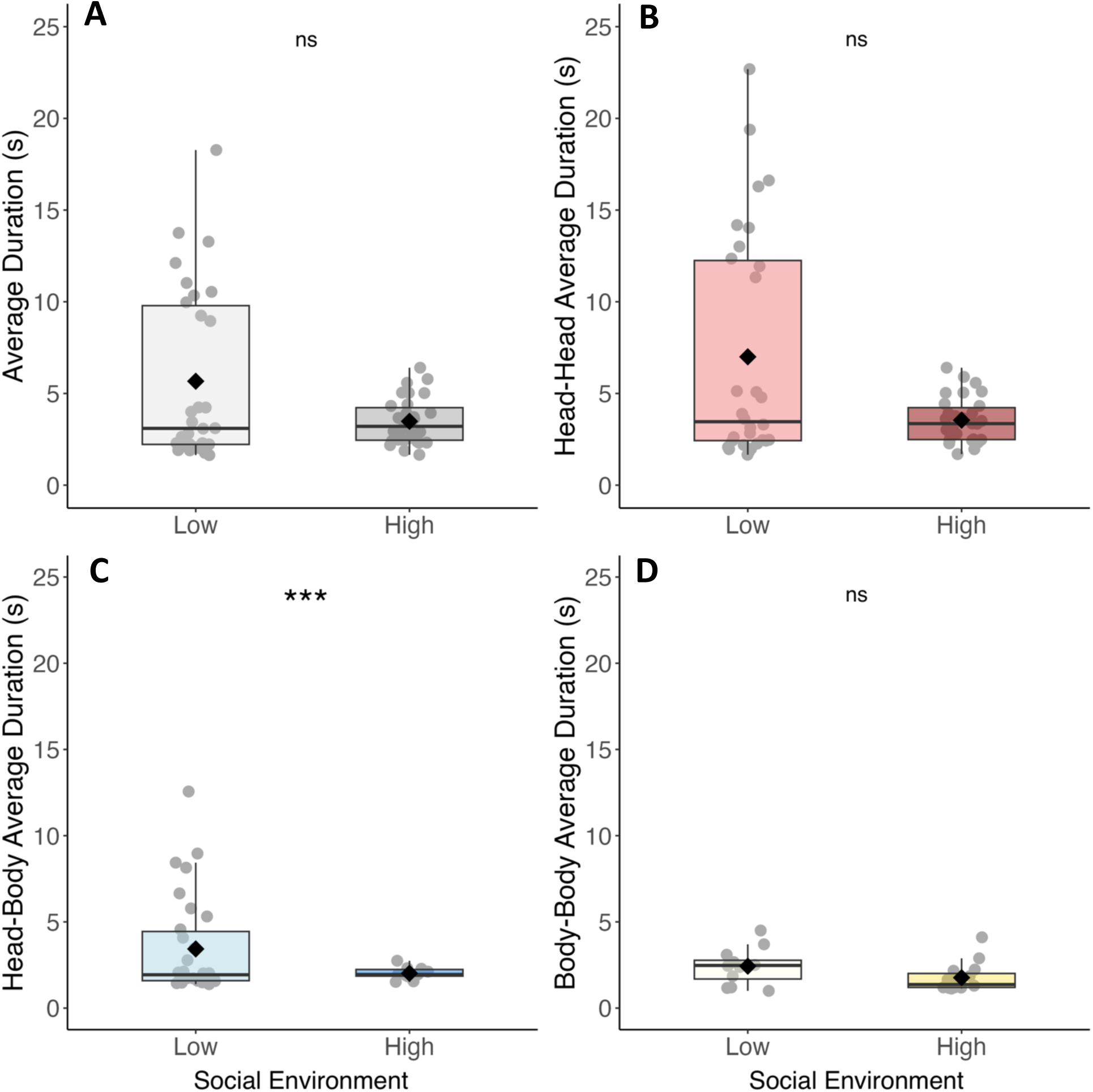
Bees in low density environments spent more time interacting. When ten bees were in larger, low-density cages, they spent more time on average having A) interactions including B) head-to-head and C) head-to-body interactions compared to when they were in smaller, high-density environments. D) There was no difference in the body-body interactions. Interactions are characterized as a boundary around the body of the individual bee overlapping. Head-to-head interactions occur when a diamond shaped boundary over the head of the bee overlap with another diamond shaped boundary with another bee. Interactions are ranked: When both occur, head-to-head interactions are prioritized. Each dot is the average for each individual in a video. Boxes represent 25-75 quartiles, whiskers indicate 95% of the data. Thick bars are medians and diamonds are means of all the data.

The performance of collective behaviors relies on effective communication of information between individuals within the group [5]. Information can be communicated indirectly or directly [5,7], but direct communication in particular offers an opportunity for information synthesis, which may then enhance the robustness and resilience of the response to a changing environment. Here, we provide further evidence that the collective fanning response is likely dependent on two types of information: thermal information and social information. Social information is modified by density in many biological systems, such as cancer metastesis, migration, and misinformation [13,51–53] and unsurprisingly has an impact on collective behaviors in social insects. In summary, our work here establishes a tractable system in which to test important hypotheses about how collective animal groups utilize environmental information to respond effectively to a changing environment.

## Acknowledgements

We thank David Farynk for his assistance in setting up and troubleshooting ABCTracker, Dr. Daniel Charbonneau for assistance with the tracking data analysis and R Code. Thank you to the Cook Lab for feedback the manuscript. Funding support for these projects came from National Science Foundation Dissertation Improvement Grant (#1406773) and Integrative Organismal Systems (#1406773).

