## Supplemental Materials for "Physical Interactions Drive Collective Thermoregulatory Behavior in Honey Bees"

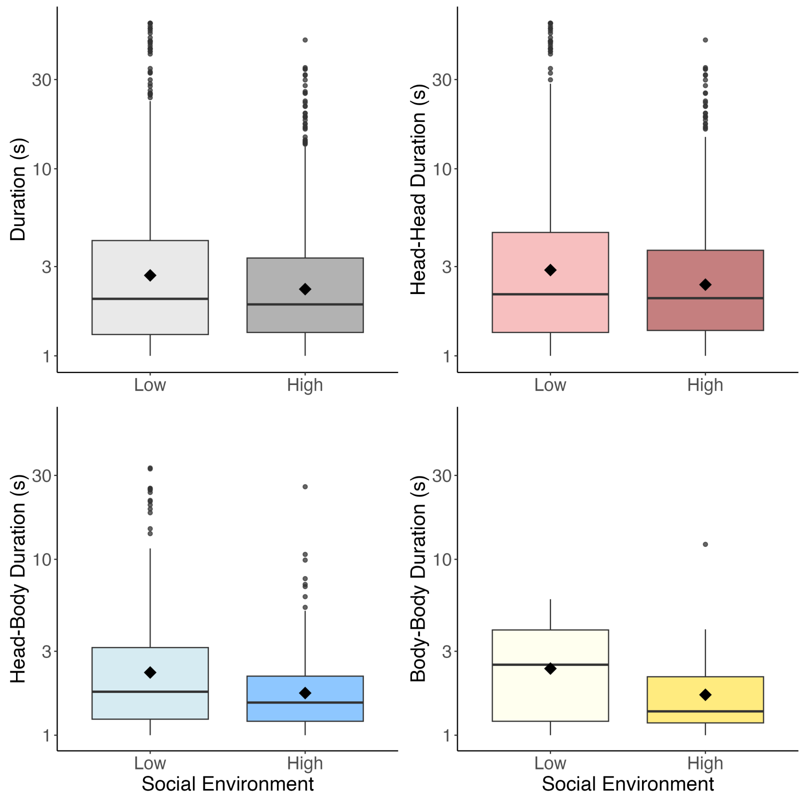


*Figure S1: Interaction duration by type, y-axis scale log(10).Boxes represent 25-75 quartiles, whiskers indicate 95% of the data. Thick bars are medians and diamonds are means of all the data.*

***Table S1A***

*Shapiro-Wilk Normality Test: Individual Number of Interactions*

| Interaction Type | Social Environment | *p*-value |
|---|---|---|
| All Interaction Types | Low Density | 0.0224 |
| | High Density | 0.1555 |
| Head-Head | Low Density | 0.5248 |
| | High Density | 0.0394 |
| Head-Body | Low Density | 0.1827 |
| | High Density | <0.0001 |
| Body-Body | Low Density | <0.0001 |
| | High Density | <0.0001 |

***Table S1B***

*Shapiro-Wilk Normality Test: Individual Average Interaction Duration*

| Interaction Type | Social Environment | *p*-value |
|---|---|---|
| All Interaction Types | Low Density | <0.0001 |
| | High Density | 0.0693 |
| Head-Head | Low Density | <0.0001 |
| | High Density | 0.1016 |

| | | |
|---|---|---|
| Head-Body | Low Density | <0.0001 |
| | High Density | <0.0001 |
| Body-Body | Low Density | <0.0001 |
| | High Density | <0.0001 |

***Table S2***

*Distribution Summary of Individual Number of Interactions*

| Interaction Type | Social Environment | *n* | *Min* | *Max* | *Mean* | *Std. Dev* | *Pctl. 25* | *Pctl. 75* |
|---|---|---|---|---|---|---|---|---|
| All Interaction Types | Low Density | 30 | 23 | 95 | 57.1 | 20.9 | 42.2 | 72.8 |
| | High Density | 30 | 30 | 351 | 157 | 96.0 | 45.5 | 237 |
| Head-Head | Low Density | 30 | 12 | 72 | 40.9 | 16.1 | 30.0 | 49.0 |
| | High Density | 30 | 17 | 176 | 76.3 | 36.6 | 46.5 | 104 |
| Head-Body | Low Density | 30 | 4 | 32 | 15.2 | 7.09 | 10.2 | 18.8 |
| | High Density | 30 | 0 | 63 | 14.3 | 21.6 | 0 | 32.2 |
| Body-Body | Low Density | 30 | 0 | 8 | 0.93 | 1.68 | 0 | 1.75 |
| | High Density | 30 | 0 | 135 | 12.7 | 34.6 | 0 | 2.75 |

***Table S3***

*Total Interaction Number Distribution by Social Environment*

| Interaction Type | Social Environment | *n* | *Mean* | *Std. Dev* | *Pctl. 25* | *Pctl. 75* |
|---|---|---|---|---|---|---|
| All Interaction Types | Low Density | 1712 | 5.30 | 9.99 | 1.30 | 4.13 |
| | High Density | 4098 | 3.14 | 3.88 | 1.33 | 3.33 |
| Head-Head | Low Density | 1228 | 5.95 | 11.2 | 1.33 | 4.57 |
| | High Density | 3286 | 3.36 | 4.12 | 1.37 | 3.67 |
| Head-Body | Low Density | 456 | 3.69 | 5.48 | 1.23 | 3.15 |
| | High Density | 430 | 2.07 | 2.12 | 1.20 | 2.17 |
| Body-Body | Low Density | 28 | 2.76 | 1.42 | 1.20 | 3.97 |
| | High Density | 382 | 2.15 | 2.30 | 1.18 | 2.15 |

***Table S4***

*Wilcoxon Test: Individual Number of Interactions*

| Interaction Type | *n* | | *W* | *p-value* |
|---|---|---|---|---|
| | Low Density | High Density | | |
| All Interaction Types | 30 | 30 | 211 | 0.0004 |
| Head-Head | 30 | 30 | 141 | <0.0001 |
| Head-Body | 30 | 30 | 606 | 0.0191 |
| Body-Body | 30 | 30 | 374 | 0.2230 |

***Table S5***

*Number of bees with recorded interactions by type*

| Social Environment | Trial | Number of bees | | |
|---|---|---|---|---|
| | | Head-Head | Head-Body | Body-Body |
| High Density | A | 10 | 8 | 10 |
| | B | 10 | 2 | 3 |
| | C | 10 | 0 | 2 |
| Low-Density | D | 10 | 10 | 7 |
| | E | 10 | 10 | 0 |
| | F | 10 | 10 | 5 |

***Table S6***

*Distribution Summary of Individual Average Interaction Duration (S)*

| Interaction Type | Social Environment | *n* | *Min* | *Max* | *Mean* | *Std. Dev* | *Pctl. 25* | *Pctl. 75* |
|---|---|---|---|---|---|---|---|---|
| All Interaction Types | Low Density | 30 | 1.64 | 18.3 | 5.67 | 4.69 | 2.22 | 9.79 |
| | High Density | 30 | 1.66 | 6.40 | 3.48 | 1.24 | 2.45 | 4.23 |
| Head-Head | Low Density | 30 | 1.66 | 22.7 | 7.00 | 6.28 | 2.43 | 12.3 |
| | High Density | 30 | 1.69 | 6.40 | 3.54 | 1.22 | 2.48 | 4.23 |
| Head-Body | Low Density | 30 | 1.39 | 2.88 | 3.43 | 2.88 | 1.59 | 12.6 |
| | High Density | 10 | 1.52 | 2.75 | 2.02 | 0.37 | 1.86 | 2.75 |
| Body-Body | Low Density | 12 | 1.00 | 4.50 | 2.43 | 1.04 | 1.69 | 2.77 |

| | | | | | | | |
|---|---|---|---|---|---|---|---|
| High Density | 15 | 1.13 | 4.11 | 1.77 | 0.82 | 1.20 | 2.01 |

***Table S7***

*Wilcoxon Test: Individual Average Interaction Duration*

| Interaction Type | *n* | | *W* | *p-value* |
|---|---|---|---|---|
| | Low Density | High Density | | |
| All Interaction Types | 30 | 30 | 465 | 0.8315 |
| Head-Head | 30 | 30 | 495 | 0.5106 |
| Head-Body | 30 | 10 | 757 | <0.0001 |
| Body-Body | 12 | 15 | 439 | 0.8650 |
